# A displacement and velocity based dual model of saccadic eye movements best explains kinematic variability

**DOI:** 10.1101/2020.03.19.998419

**Authors:** V Varsha, Radhakant Padhi, Aditya Murthy

## Abstract

Noise is a ubiquitous component of motor systems which leads to behavioral variability of all types of movements, including saccadic eye movements. Nonetheless, systems-based models of saccadic eye movements are deterministic and do not explain the observed saccade variability, only their central tendencies. Using stochastic models, we studied the variability in saccade behavior to test and distinguish between previously proposed deterministic saccade models. For this, the inter-trial variability in saccade displacement trajectories of human subjects was quantified while they performed repeated saccadic eye movements to a peripheral target. Based on fits to the data, we showed that existing models based on either displacement or velocity failed to capture the observed patterns in the variability of saccade trajectories. However, the observed behavior was captured by a dual control system, using a combination of displacement and velocity signal. The proposed model fits the mean displacement trajectory as well as the existing deterministic models. Taken together, our results suggest that the saccade system uses both desired displacement and velocity information.

**New and Noteworthy:** We studied saccade behavior with a focus on the variability of the saccade trajectory. A stochastic model of the saccade system suggests that a dual control involving the control of displacement and velocity explains saccade behavior better than previously proposed models that utilize only displacement or velocity information. Our study resolves previous ambiguity regarding the use of displacement or velocity signals to guide saccades and provides a natural explanation for neural recordings that indicate multiplexing of displacement and velocity related information in the firing activity of neurons in the superior colliculus, a critical node in the oculomotor network that codes for saccadic eye movements.

---

The motor system is inherently noisy and the consequence of this can be observed in behavior in the form of motor variability (1). Variability is not only observed between subjects but also within a subject when they repeatedly perform the same movement. If such intra-subject variability is an intrinsic characteristic of the movement generating system itself, a study of such variability should provide insights into how movements are generated. For example, signal-dependent noise has been used before to model the stochastic properties of the saccade system but its applicability has been restricted to formal models of optimal control (2–5). It has not been used in the more standard control systems models that have motivated much of the research on saccadic motor control (except for (6)). In contrast, much of the control systems-based modeling of saccades that have been used to explain the main sequence of saccades, are deterministic and ignore their variability (7–10). Thus, it remains unknown whether these models provide an adequate account of the variability in saccade kinematics. In the internal feedback model of saccade generation, the controller gets the desired input which is used to calculate the motor error with the help of a local internal feedback signal. This motor error is converted to a pulse output and is integrated to generate the pulse step signal required for saccade generation (7). Earlier models considered the desired input to the saccade system to be final position (11) and later models suggested that it was the final displacement (8, 10). Recent evidence, however, reveals that the desired input to the system could be a dynamic signal of instantaneous displacement or velocity (9, 12). Deterministic models based on static signals like final displacement (8, 10) as well as dynamic signals like instantaneous displacement (9), however, have similar predictions. Thus, it remains an open question, whether the saccade execution system is based on just the final displacement or a dynamic signal like velocity. In this study, we show that a computational model of saccade generation system which uses both velocity and displacement signals to generate and control saccades, explains the variability in saccade trajectories.

## METHODS

### Experimental details

#### Subjects and set up

Experiment was performed on 20 subjects (equal number of male and female) from the age group of 21 − 33 years. Participants had normal vision or corrected to normal vision. The protocol was approved by the Institutional Human Ethics Committee and all the participants gave their informed consent. During the experiment, participants sat on a chair in front of a monitor, with their heads locked at the temples and their chins resting on a chin-rest to minimize head movements. The stimulus was displayed on a monitor kept at 57 cm in front of them. This ensured that 1 cm on the screen subtended a visual angle of 1°. Videosync software along with Tempo software (Reflective Computing, USA) allowed the display of the stimulus and control of the behavioral task. The eye position was recorded in real-time at 240 Hz. The eye tracker used was a monocular infrared pupil tracker (ETL-200, ISCAN) placed in front of the monitor and focused on the eye to be tracked.

#### Task Design

The task started with a standardization block in which the subject was made to fixate on all the target positions to calibrate the eye tracker data. The task consisted of an average of 92 blocks of 15 trials each presented to each subject. Each trial started with a white fixation box of size 0.35^°^ appearing at the centre of the screen. The subject had to fixate on this white box for 400 ms with a jitter-time of maximum 160 ms. The fixation box kept toggling in case the subject did not fixate properly. The toggling increased the saliency of the stimulus and facilitated fixation of the eye on it. A correct fixation required them to be in an electronic window of 4.7°, centred at the fixation box, which was not revealed to the participants. A green target of size 0.7^°^ appeared at the periphery at one of 15 possible target positions following successful fixation. Immediately after the appearance of the target, a saccadic eye movement was to be made. If the subject did not move their eye until 400 ms, the trial was aborted. The saccadic eye movement had to be completed within 100 ms. If the subject made a successful eye movement within this time, a green tick mark and an auditory tone audio feedback were given. A red cross mark was shown at the end of the trial in case the subject did not complete the movement within the allowed reaction time and saccade movement time. Saccades that foveated the target within an electronic window of 2.9^°^ were considered as successful and were denoted by the success tone. In this work, we have only analysed 12^°^ horizontal saccades made to the left as horizontal saccades involve only a single muscle (either the medial or lateral rectus); in contrast, vertical and oblique saccades involve multiple extraocular muscles (the superior or inferior recti as well as the superior or inferior oblique), that make their contributions to kinematic variability harder to understand.

### Data analysis

#### Saccade detection

The data was first passed through a convolution filter which averaged each data point with its two immediate neighbouring points to remove unwanted jitter. The velocity of each saccade in each trial was obtained by computing the differential between successive position data. The start of the saccade was marked when the velocity crossed 30deg/s and the end of saccade when the velocity fell below this threshold. Saccades with blinks are omitted by checking if the velocity crossed 700deg/s. Only the first saccade was processed further.

#### Pre-processing

A set of pre-processing steps were carried out for strict outlier removal. This was followed by a time normalization. Saccades made to the same target could have different duration as can be seen in Figure 1. Hence, time normalization was required for obtaining an equal number of bins of data in each trial across which the mean and variance could be calculated. These processing steps are discussed in detail in the supplementary material (S1).

**Fig. 1.**
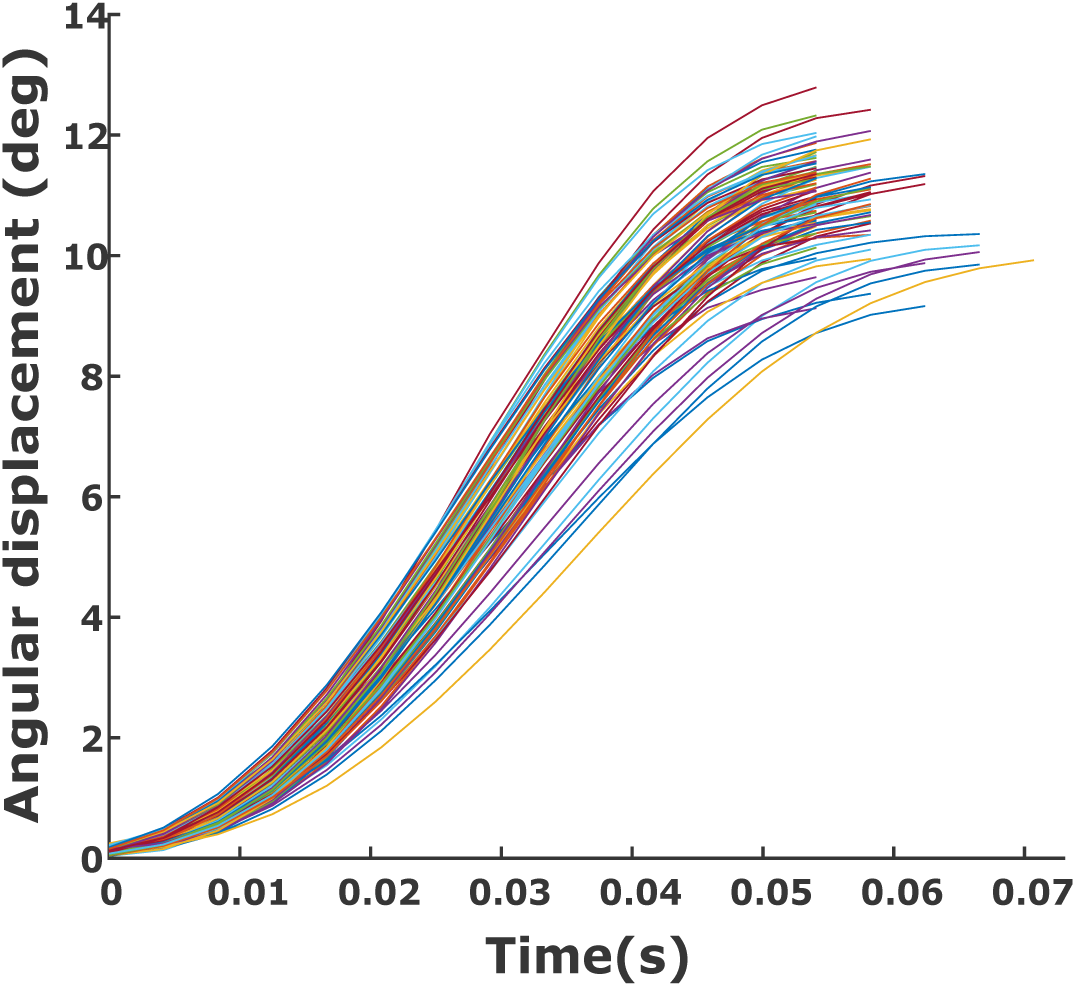
Variability in angular displacement of saccades. Each line in the plot is angular displacement data from a trial of saccade task.

#### Mean and variance

The mean of the saccade angular displacement trajectory and inter-trial variance was obtained for each subject individually. The mean 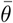 was calculated as the sample mean of *N* trials at each time bin *t*_*k*_ and is given by the formula

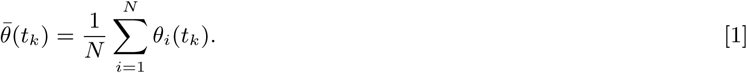

The inter-trial variance *θ*_var_ was calculated from the sample variance at each time bin using the same N trials by the formula

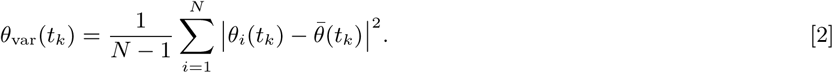

The number of trials used (N) was chosen as 50. This was based on the observation that the change in variance profile was less than 0.1deg^_2_^ after 40 trials and that profiles remained qualitatively the same if 50 trials were randomly sampled from a larger population of trials.

#### Statistical tests

Statistical tests were used for comparison between experimental observations and model predictions. All data were checked for normality using the Lilliefors test. If the data were normally distributed, then a two-tailed t-test was used, otherwise, a Wilcoxon signed-rank test was used for testing statistical comparisons. Statistical significance is marked on the figures according to the following norm: *p* < 0.05 and *p* > 0.01 : *, *p* < 0.01 and *p* > 0.001 : **, *p* < 0.001 : ***. The p-value is mentioned when it is greater than 0.001.

#### Fit error quantification

The error *e*_*k*_ in the data fit is quantified as a normalized error between the prediction of the model 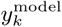 and experimental data 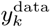 for all *m* time bins and is defined as

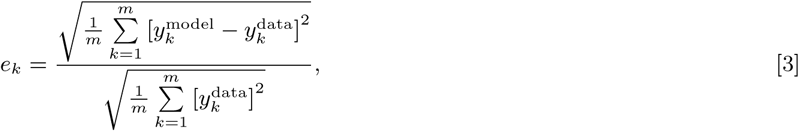

for any output variable of interest *y*, which can be mean angular displacement or displacement variance.

### Model of Saccade Generation

The dual control model proposed for a stochastic saccade generation system consists of four major parts. A displacement block, a velocity block, a pulse step generator, and the oculomotor plant (Figure 2). The stochastic characteristic of the model is due to the presence of noise that affects the system at multiple levels: planning noise that affects the input, the burst neuron noise that acts on the burst generator, and the oculomotor noise at the input to the oculomotor plant (6).

**Fig. 2.**
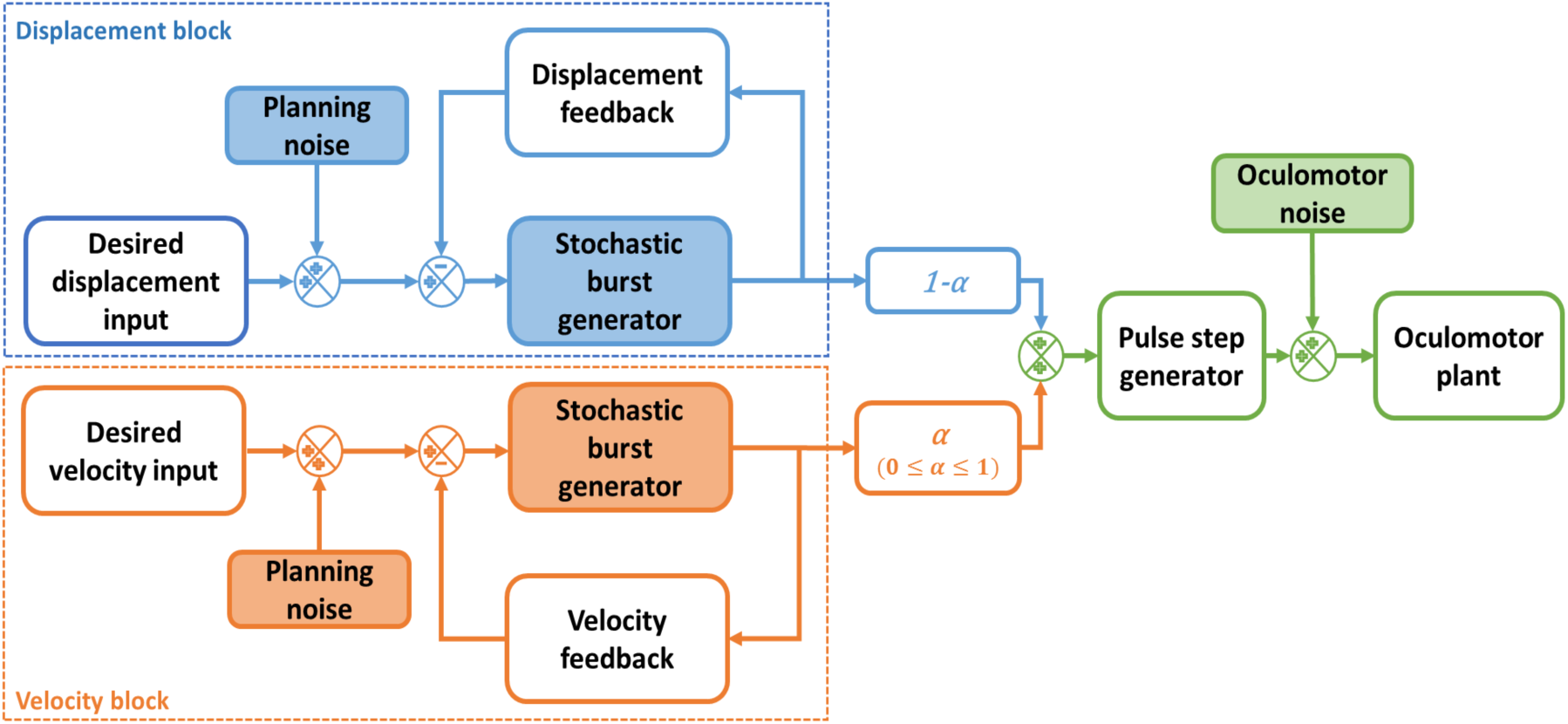
Schematic of the stochastic dual model. The displacement block and the velocity block have three components: the desired input with planning noise, the stochastic burst generator, and the internal feedback. A convex combination of the output of these blocks is given as input to the pulse step generator. The pulse-step generator output is added with oculomotor noise and becomes the input to the oculomotor plant. The arrows represent the direction of the flow of information. Shaded components are either noise generators or include noise generators within.

#### Displacement block (see Figure 2)

The displacement block includes a stochastic burst generator and an internal feedback loop. It receives the desired input which is affected by planning noise. The desired input *r*_*d*_(*t*_*k*_) is modelled as a step function of amplitude *A*_*d*_ acting only for the mean saccade duration *t*_*f*_.

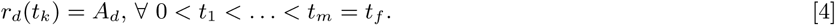

This signal is assumed to encode the population firing rate of neurons in the superior colliculus that fire for a fixed saccade amplitude. The planning noise *n*_*pnd*_(*t*_*k*_) is added to the input before it is compared with the feedback signal. It is modelled as white noise *w*_*pnd*_(*t*_*k*_) that is scaled by a gain *k*_*pnd*_ for all time bins *t*_*k*_. The noise distribution changes only with a change in saccade amplitude. The noisy desired input *r*_*nd*_(*t*_*k*_) becomes

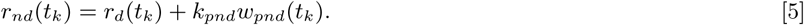

The input along with the output of the internal feedback loop is used to estimate the instantaneous motor error *z*_*d*_(*t*_*k*_). The internal feedback loop in the case of the displacement block consists of an integrator that converts the output of the burst generator, which encodes velocity, to an instantaneous displacement. The burst generator converts the motor error *z*_*d*_(*t*_*k*_) to a pulse that is required to overcome the inertia of the eye and is modelled as a saturating function of motor error (13). This signal is hypothesized to be affected by additive signal-dependent noise (14) called burst neuron noise *n*_*bnd*_(*t*_*k*_) = *k*_*bnd*_*b*_*d*_(*t*_*k*_)*w*_*bnd*_(*t*_*k*_). Thus, the output of the stochastic burst generator is given by

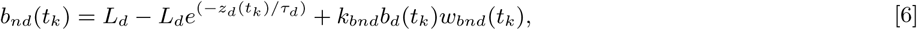

where 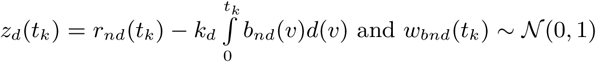 and independent for all 0 < *t*_1_ < … < *t*_*m*_ = *t*_*f*_. Note that, *b*_*d*_(*t*_*k*_) is the burst generator output without noise, *L*_*d*_ is the gain of the burst generator block, *k*_*d*_ is the internal feedback gain of displacement block and 𝒩 (0, 1) represents a Gaussian distribution with mean 0 and variance 1. The noisy output *b*_*nd*_(*t*_*k*_) is the input to the internal feedback loop.

#### Velocity block (see Figure 2)

Like the displacement block, the velocity block also consists of a burst generator and an internal feedback loop. The major difference is that the information at the input is encoding the desired velocity profile, unlike the desired displacement profile. Hence, the input *r*_*v*_(*t*_*k*_) is a dynamically varying signal and is modelled as a Gaussian function given by

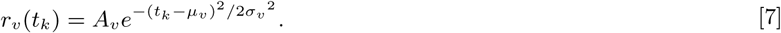

In Eq.7, *A*_*v*_, *µ*_*v*_ and *σ*_*v*_ are free parameters that are estimated. This input *r*_*v*_(*t*_*k*_) is corrupted by a planning noise *n*_*pnv*_(*t*_*k*_) which is signal-dependent. The planning noise in this case is modelled as white noise *w*_*pnv*_(*t*_*k*_) that is scaled by a gain *k*_*pnv*_ and the input *r*_*v*_(*t*_*k*_) for all time bins *t*_*k*_. The noisy input *r*_*nv*_(*t*_*k*_) is given by

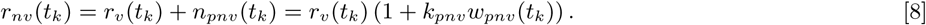

The noisy input is used along with the output of the internal feedback signal to obtain the motor error *z*_*v*_(*t*_*k*_) that goes into the burst generator. Since the input is velocity, the internal feedback is just a proportionally weighted burst generator output. Hence, the burst generator is also modelled as a linear transformation with gain *L*_*v*_ as in (9). The noisy burst generator output *b*_*nv*_(*t*_*k*_) after incorporating a signal-dependent burst neuron noise is modelled as

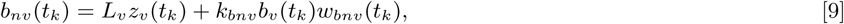

where *z*_*v*_(*t*_*k*_) = *r*_*nv*_(*t*_*k*_) − *k*_*v*_*b*_*nv*_(*t*_*k*_) with *k*_*v*_ being the internal feedback gain of velocity block.Note that, *b*_*v*_(*t*_*k*_) is the burst generator output without noise in the velocity block and the white noise *w*_*bnv*_(*t*_*k*_) ∼ 𝒩 (0, 1) is independent for all 0 < *t*_1_ < … < *t*_*m*_ = *t*_*f*_. The basic philosophy of the deterministic part of the displacement block is similar to (10) and that of the velocity block is similar to (9).

#### Burst to saccade displacement conversion

The proposed dual model uses a combination of displacement and velocity information. This multiplexing is assumed to be carried out at the output of the burst generators and is modelled as a convex combination of the burst generator outputs of the two blocks. This can be written as

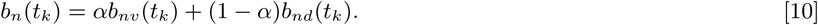

This burst is converted into a pulse and step by the pulse step generator to generate the control signal required by the oculomotor plant. The noisy control signal with signal-dependent oculomotor noise is given by

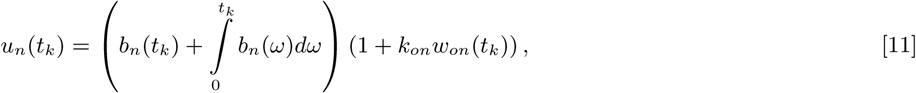

where *w*_*on*_(*t*_*k*_) ∼ 𝒩(0, 1) and independent for all 0 < *t*_1_ < … < *t*_*m*_ = *t*_*f*_. This control signal, when given to the oculomotor plant, generates the saccadic displacement. The plant is modelled as a second-order spring mass damper system with an impulse response *p*(*t*_*k*_) and then the angular displacement can be obtained as

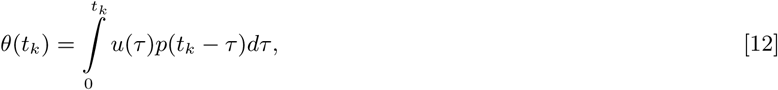

where 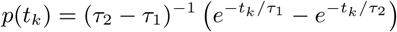. The values of the time constants are taken as *τ*_1_ = 223 ms and *τ*_2_ = 14 ms as used in (2). These equations can be expanded by using definitions of mean and variance to get the mean angular displacement and variance in angular displacement predictions of the model in terms of the free parameters in the model. The free parameters are estimated such that the error of these predictions with the experimental mean and variance is minimized. The details of these derivations are discussed in the supplementary material (S2).

## RESULTS

To test whether the type of information used for saccade control is driven by a displacement (8, 10) or a velocity (9, 12) signal, we designed a stochastic dual control model of the saccade generation system which utilized a convex combination of the displacement information and velocity information, as shown in Figure 2. Depending on the weightage assigned to the velocity part and the displacement part, the dual model predicted qualitatively different variance profiles: the more the weightage to the displacement model, the lower was the variance towards the end of the movement, whereas the profile switched to a monotonically increasing profile as the weightage in the velocity component increased. An example of this behavior of the model is shown in Figure 3.

**Fig. 3.**
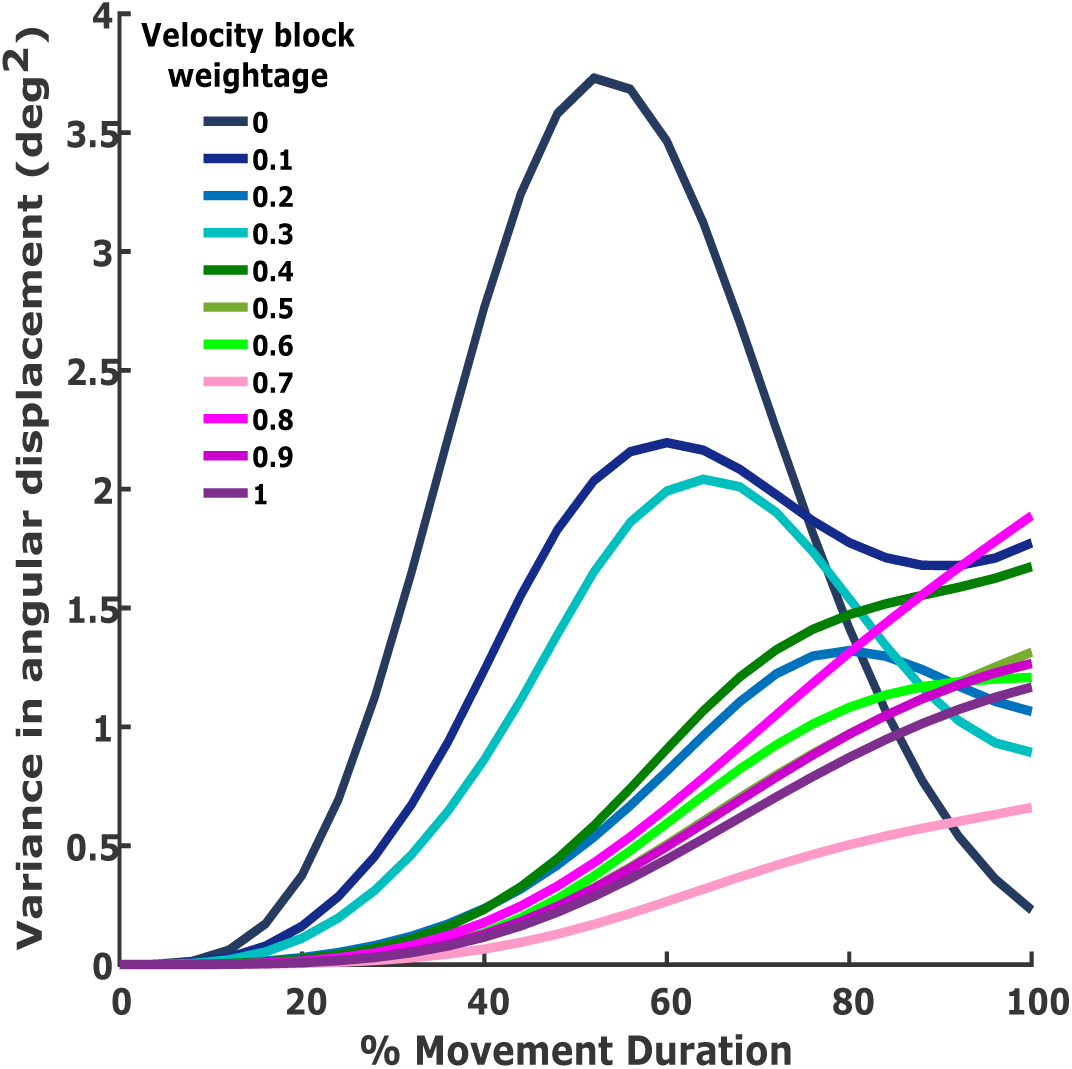
Stochastic dual model prediction. Each trace is the variance in angular displacement predicted by the stochastic dual model for a simulated example system. Note that increasing the velocity weightage changes the variance pattern from a decreasing profile towards the saccade end to a monotonically increasing profile.

To test the predictions of the model, the inter-trial variance in saccade behavior of 20 healthy human participants were calculated. The mean angular displacement profiles for all 20 participants of the experiment showed consistent mean profiles (see data shown in panel A in Figure 4) and there were small variations in mean amplitude across subjects around the target amplitude, which is consistent with previous observations. As observed in numerous other studies like (6, 10, 15, 16), inter-trial variability was observed in saccade trajectories of repeated trials of saccades made towards the same target (Figure 1). The inter-trial variance profile for the 20 subjects showed a wide variety even qualitatively (see data shown in panel B in Figure 4), as predicted by the proposed dual model (Figure 3). All profiles were monotonically increasing from the start of the saccade till the midpoint. Three qualitative types of profiles were observed based on the later part of the variance evolution. The dominant type was the profile that continued to monotonically increase until the end of the movement. Less frequent were the profiles of inter-trial variance that saturated towards the end of the saccade duration. We also observed a third type of profile, which showed a peak and decreased towards the saccade end. Although only 15% of the population showed such evident decreasing profiles, in 9 out of 20 subjects the endpoint variance was lower than the peak value in the variance profile, suggesting that this third profile was a robust pattern as well. The proposed stochastic dual model was fit to the experimental data of profiles of mean angular displacement and profiles of inter-trial variance in angular displacement. The fits are shown for all the 20 subjects (Figure 4).

**Fig. 4.**
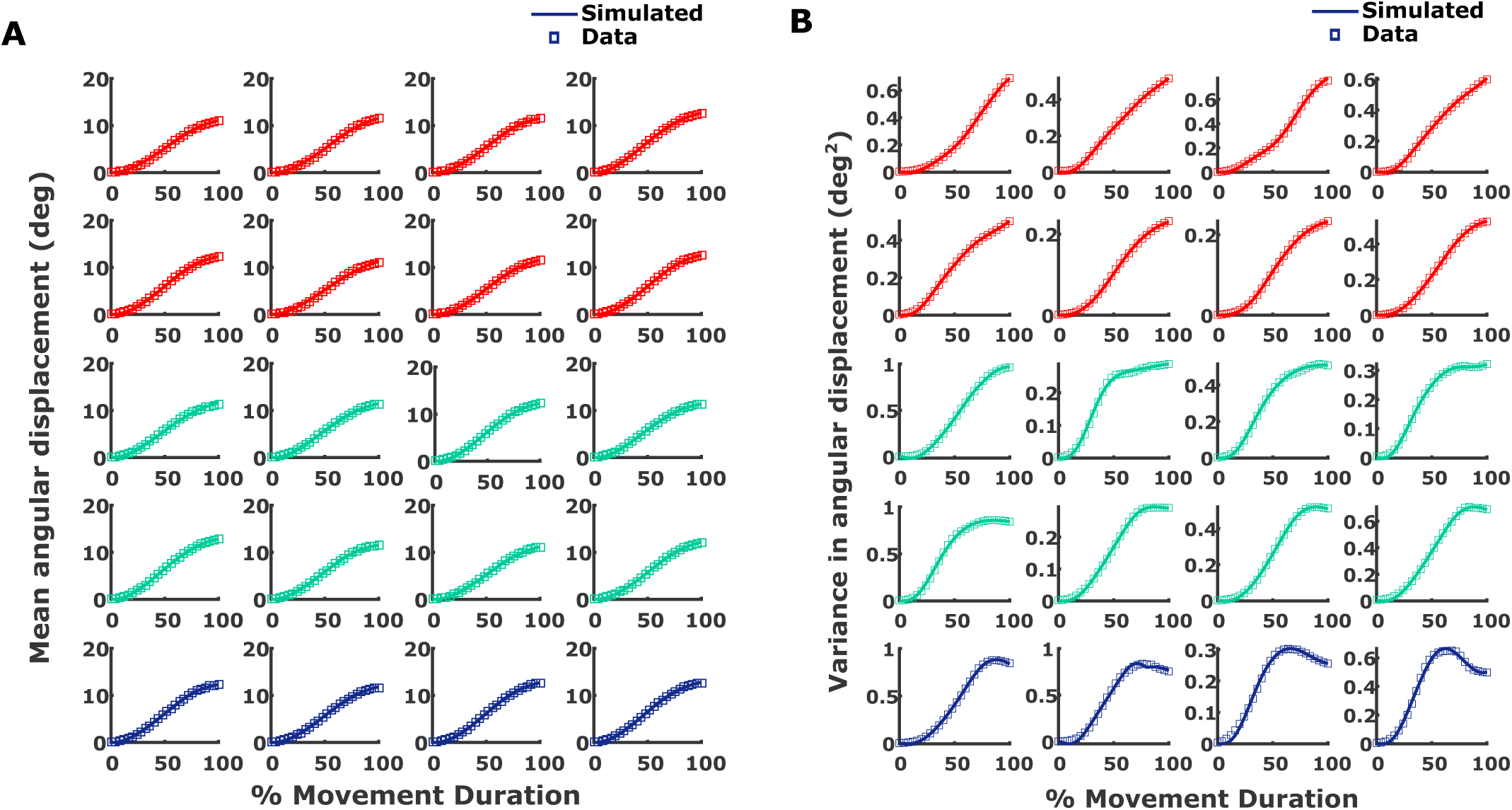
**A. Mean angular displacement fits of the stochastic dual model.** Each subplot represents a subject. The square markers are experimental data points used for fitting and the solid lines are simulated data for that subject for fitting. The first two rows in red color are for subjects with increasing variance profiles, the third and fourth rows in cyan color are subjects with saturating profiles and the last row in navy-blue color are subjects who showed decreasing variance profiles. **B. Inter-trial variance fits of the stochastic dual model**. The subplot layout and color coding are the same as A. The variance data are represented by square markers and the fit for the dual model are shown using solid lines.

The fit errors for the mean angular displacement profile of all subjects were less than 3%. The averaged fit error across 20 subjects was 1.33%. The stochastic dual model was able to fit individual variance profiles though they varied qualitatively. All fit errors were less than 3.5% and the averaged errors in the fits were 1.71% across the 20 subjects. The errors in the fit for each subject were quantified as in Eq.3, and is shown for all 20 subjects for mean angular displacement as well as displacement variance (Figure 5).

**Fig. 5.**
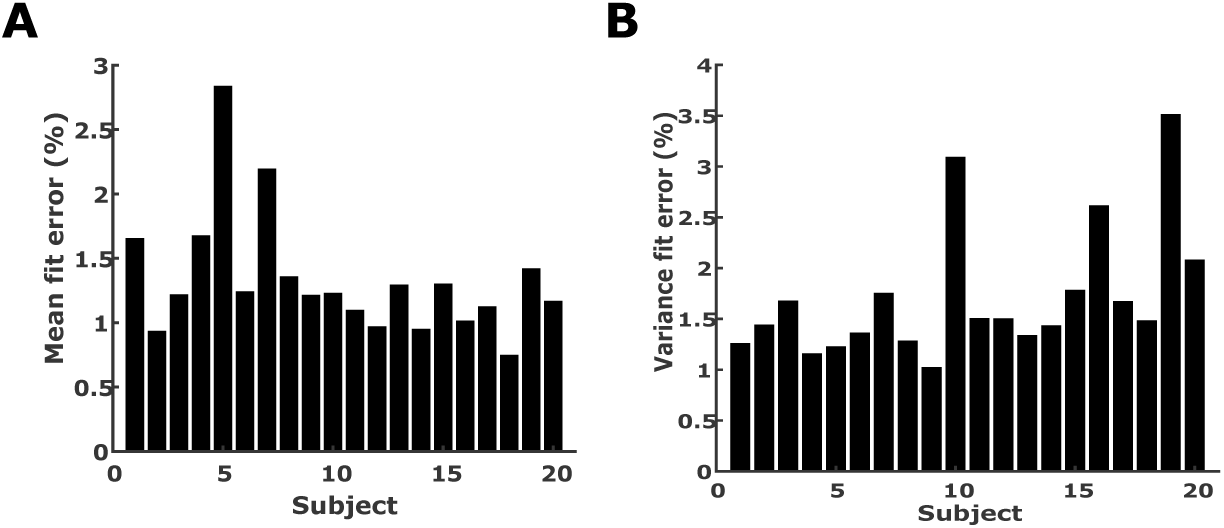
**A. Mean fit error quantification of the stochastic dual model.** The fit errors of the dual model for the mean of angular displacement for 20 subjects are shown in the bar plot. **B. Variance fit error quantification of the stochastic dual model**. The bar plot shows the fit error of inter-trial variance in angular displacement for 20 subjects.

The reduced versions of the proposed dual model which would be equivalent to existing philosophies of saccade generation based on just the displacement or velocity were also tested. For all 20 subjects, the deterministic versions of the reduced models based on displacement only and velocity only were able to predict the mean angular displacement with a fit error of less than 4%. The free parameters in the model were estimated by minimising only the error between the experimental data and the prediction of the model of the mean observed displacement. The average of fit errors of mean displacement obtained across the 20 subjects was 1.1% for the displacement model. The velocity model performance was also comparable to the displacement model in case of prediction of the mean angular displacement. The mean error in fits across 20 subjects was 1.58%.

Both the displacement and velocity models were incorporated with noise blocks and a stochastic version of the same was designed. The fit errors increased compared to the deterministic model because, in the optimization process for free parameter estimation, the error function used was a sum of the error in mean prediction and error in prediction of variance. The inter-trial variance profiles obtained from behavior were fit using both the stochastic displacement as well as the stochastic velocity model. The fit errors of displacement and velocity model for inter-trial variance profiles in angular displacement in the data, when averaged over all subjects, were 13.73% and 13.5% respectively. Additionally, both models failed to even qualitatively capture observed variance profile types. The proposed stochastic dual model performance was compared to these reduced versions of stochastic displacement and velocity model (Figure 6). For the mean angular displacement profile fits, which are obtained by considering system without noise, the displacement model and dual model had comparable fit errors (Signed rank test: *p* = 0.08). The fit errors of the stochastic velocity model and dual model were also similar (Signed rank test: *p* = 0.26). However, in the case of the variance profile prediction, the stochastic dual model performed better than the stochastic displacement model (Signed rank test: *p* < 0.001) and stochastic velocity model (Signed rank test: *p* < 0.001).

**Fig. 6.**
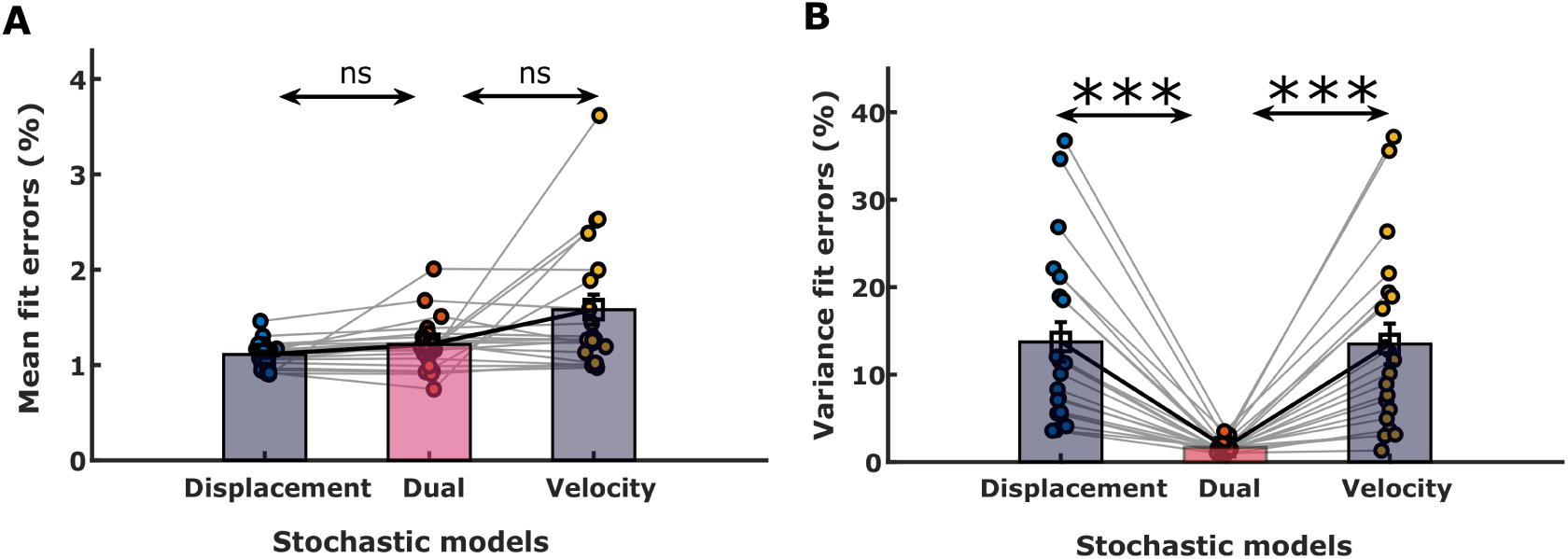
**A. Comparison of mean fit errors.** The bar plot shows a comparison of the fit errors of mean angular displacement profiles of 20 subjects for the stochastic models. The pink bar represents the dual model and the grey bars are reduced models. Circular markers represent the fit errors for each subject. Blue is the displacement model; orange is the dual model and yellow is the velocity model. The black square marker at the top middle of the bars with the cross-hairs show the mean and standard error of fit errors for all subjects. **B. Comparison of variance fit errors**. The bar plot shows a comparison of the fit errors in angular displacement variance across 20 subjects for the stochastic dual model and the reduced models. The colour coding and markers are the same as in A.

Since the dual model had a larger number of parameters, an F-test was performed to see if the fit errors of the dual model had a significant difference compared to its two reduced versions, i.e. displacement model and velocity model. The F-test showed a significant difference between the total fit errors (i.e. the sum of the mean displacement fit error and displacement variance fit error) of the stochastic dual model and stochastic displacement model in 14 out of 20 subjects. The difference was significant even when errors were averaged across all 20 subjects (F-test: *p* < 0.001). The F-test also showed a significant difference between stochastic dual model and stochastic velocity model comparisons for 18 of the 20 subjects (Table 1). Even across all the subjects, this difference was significant (F-test: *p* < 0.0001).

**Table 1.**
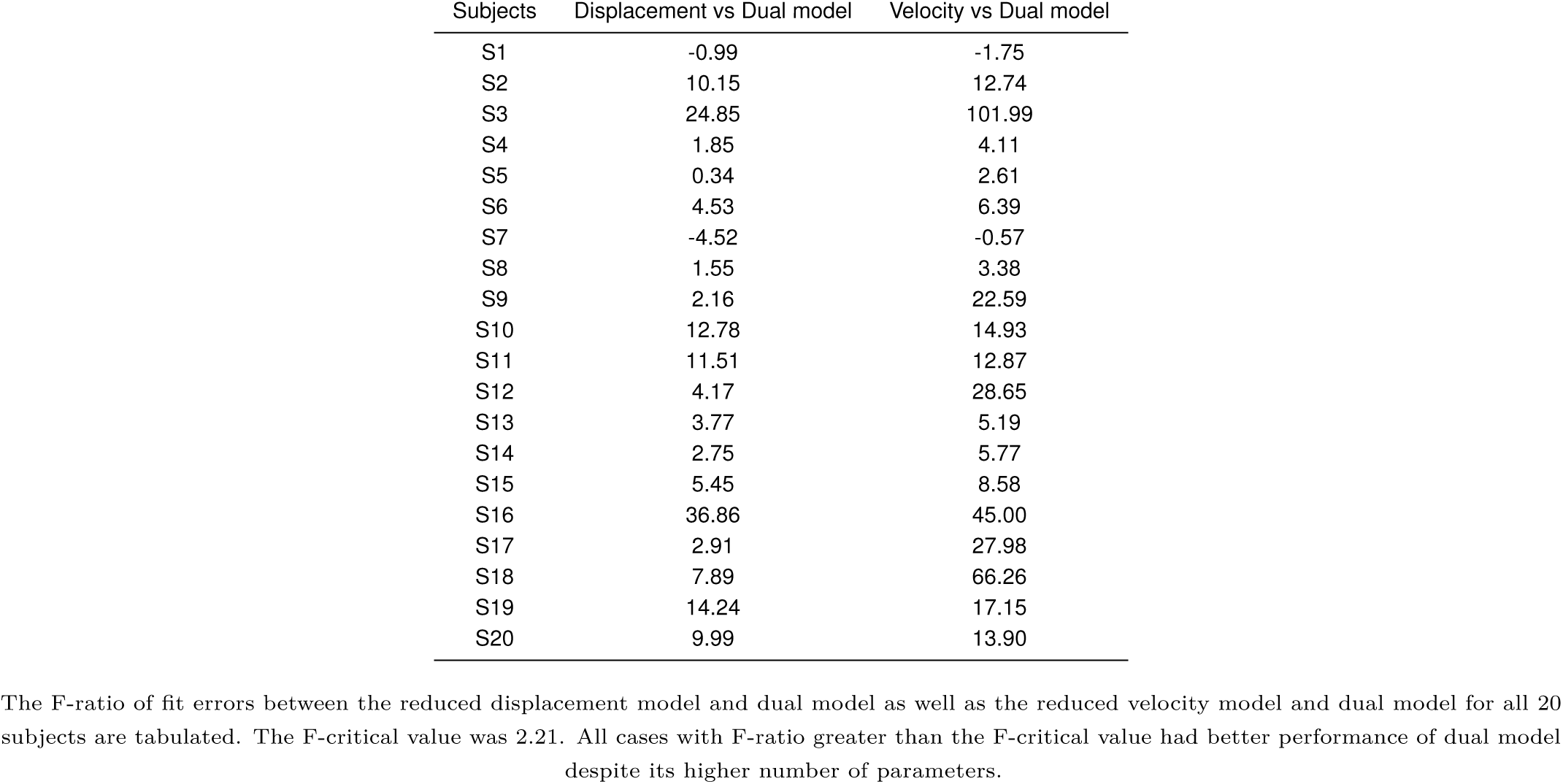
F-ratio for comparison between performance of stochastic model.

## DISCUSSION

We have quantitatively studied the variability profiles of saccade trajectories and observed that there are different types of variance profiles that can be exhibited by subjects. Using a control systems approach of modelling the stochastic saccade generation system, we suggest that saccadic eye movements use a combination of displacement and velocity information through an internal feedback loop. The proposed dual model for stochastic saccade generation fit all types of variance profiles observed in experimental data while other reduced models failed to do so.

In contrast to the well-studied main sequence of saccades, few studies have quantified variability in saccade endpoints and peak velocities (10, 15–18). Further, apart from the study by Eggert et.al. (6), no study to the best of our knowledge has incorporated saccade trajectory variability into existing models. However, unlike the current study, whose primary objective was to distinguish between velocity-based and displacement-based models, the primary purpose of their work (6) was to test the contribution of different sources of noise to the observed variability. A major conclusion of their study was that all sources of noise, such as signal-dependent planning noise, signal-dependent oculomotor noise, and signal-dependent premotor noise entering within an internal feedback loop, were necessary to explain the variance data. This forms the basis of our stochastic model and hence, our proposed model also incorporated three sources of noise as suggested in (6) with one major difference being that our dual model had an extra signal-dependent planning noise affecting the desired velocity input and a separate signal-dependent premotor noise entering the internal feedback loop in the velocity block (refer Figure 2). Furthermore, like the study of Eggert et al. (6), we did not consider constant noise or temporal noise that is proposed in (15). This may not be required in our case as we fit the model to data from single amplitude saccades, where the mean duration and mean amplitude is assumed to be their optimal values.

Like previous work (2, 3, 6), the basic motivation behind considering signal-dependent noises were the studies which showed that the motor planning is determined by the signal-dependent nature of the noise and provided empirical evidence for the same in muscles (14, 19). They had shown that the source of this noise is not neuromuscular in nature but reflects the noise in the motor command at the premotor neuron level. Taken together with the evidence provided in (9) that the noise in the superior colliculus neurons is signal-dependent, the partitioning of the motor noise into constituent components, as has been done here and by Eggert et. al. (6), seems to be a valid modeling strategy. The signal-dependent noise in the model are assumed to have a Gaussian distribution with the variance scaled by the square of the expectation of the signal. However, given that the scaling factors are free parameters that are tuned to the saccade data, the results would be invariant to the assumption of whether the noise variance scales linearly (as in Poisson statistics) or quadratically. Here, we use the best fit of respective models to the saccade data, irrespective of the magnitude of the different sources of noise, to compare between displacement, velocity, and the proposed dual model.

We show that the saccade generation system, in the presence of noise, is best explained by a convex combination of displacement and velocity information. The displacement-only model predicts a variance profile that has a peak and then decreases towards the end of the saccade. The velocity-only model, which can be considered similar to the instantaneous displacement control model (9), predicts monotonically increasing variance profiles. These increasing profiles of variance were also the commonly observed pattern in (6). However, contradictory to our observation from the dual model, in their work on monkey saccade data, displacement only model was used to fit the increasing and saturating variance profiles. This may be because, in their model, the internal control signals at the oculomotor neurons and premotor burst neurons were generated using the main sequence data. This could have led to an implicit incorporation of the desired velocity input like characteristics into the model. It is worth noting that the proposed dual model is a more generalised philosophy and can be reduced to displacement only or velocity only model by setting the weightage parameter appropriately (*α* = 1 leading to velocity only model and *α* = 0 leading to displacement only model). Nevertheless, both reduced models fail to capture the variety of variance profiles observed in our data, whereas these profiles can be predicted by a convex combination of the displacement and velocity blocks (refer Figure 6). This is clear from our observation that the weightage parameter *α* has a distribution with a mean 0.39 and a standard deviation 0.3 across different participants.

A natural question to be asked is the basis for qualitative differences in the profiles of inter-trial variability across subjects. Although we do not know the answer to this question, the newly proposed dual model for stochastic saccade generation system suggests that saccades make use of a multiplexed representation. It is consistent with studies that have reported the average firing rates in collicular cells, that provide input to the brainstem saccade generation system, to be correlated with dynamic signals like saccade velocity or instantaneous displacement (9, 12), or the peak saccade velocity (20–23). It is also possible that such information is multiplexed as a complex representation as a consequence of learning such representations in neural networks, as suggested in (24). In any case, having a multiplexed system with both displacement and velocity information seems to be a more robust solution for oculomotor control of fast saccadic eye movements in the absence of sensory feedback.

## ACKNOWLEDGMENTS

We acknowledge the funding agencies which supported this study. The experiments were supported by grants by a Department of Biotechnology-Indian Institute of Science (DBT-IISc) partnership programme grant. Varsha Vasudevan was supported by a fellowship from Indian Institute of Science.

No conflicts of interest, financial or otherwise, exists for this work.

## SUPPLEMENTARY MATERIAL (S1)

### Pre-processing of saccade data

All saccades made by the subject irrespective of if they were given a feedback for being correct or not during the task were considered in the analysis. Saccades outside three standard deviations of the saccade amplitude distribution for each subject were eliminated as outliers. The reduction in trials due to this criterion was only 1.54% on average, excluding two subjects who showed a slightly higher percentage reduction of 34.65% (*S*17) and 11.69% (*S*18). This was attributed to the absence of post saccade time in the experiment design, which in these subjects, caused the eye to drift out of the target window immediately after reaching the target or a tendency to make hypometric saccades accompanied by a small corrective saccade to reach the target. Additionally, although the fixation box size was 0.35°, subjects spanned almost the entire fixation window of 4.7°, although they were unaware of this freedom allowed in the experiment. This caused an average fixation standard deviation of 2.4° in the horizontal direction and 2.3° in the vertical direction across subjects. There could be multiple possible reasons for this fixation jitter. The fovea has a size of 1°, thus contributing to some of this deviation. Further small drifts, which cannot be controlled by the subjects, could be an added factor. This initial variability was corrected by a normalization procedure that subtracted the mean position data from 20 ms before the start of each saccade from the entire trajectory for each trial (similar to the procedure in (15, 25)). This normalization procedure was performed for the horizontal and vertical position data separately, and then the angular displacement was calculated.

A set of secondary checks were carried out on the marked saccades for further outlier elimination. Only trials that did not have any sudden changes in the trajectory were included. This was done by excluding all trials in which at any time point during a saccade, the difference between consecutive values was greater than half of the standard deviation of the values taken by the saccade within that trial. Trials with saccade durations beyond three standard deviations of the mean duration distribution were eliminated.The final step in the pre-processing after outlier removal was time-normalization. All saccades from repeated trials to the same target have measurably different durations. An example of the same can be seen in Figure 1 in main text and this is due to the different trajectories the subject take while repeatedly making saccades to the same target. Thus, to compare across trials, saccades were re-binned in time from the start of the saccade to its end. For this, the data points of horizontal and vertical eye position from 20 ms before to 20 ms after the saccade was fit using a seventh order polynomial fit. The seventh order polynomial ensured that the acceleration was fit with a fifth-order polynomial, which gave good acceleration profiles with clear acceleration and decceleration peaks. This polynomial function obtained by fitting the angular displacement data of saccade was differentiated to obtain the velocity of the saccade. Using this velocity data, the start and end of the saccade were readjusted by taking start as the time when the velocity increased above 10% of the peak velocity in that trial and the end of saccade as the time when saccade velocity fell below this threshold. Trials with double peaks in their velocity profiles or peak velocity outside three standard deviations of the peak velocity distribution of the subject were eliminated from further analysis. The readjusted saccades between the new start and end time were calculated using interpolation. The saccade start was considered completion of 0% of the final duration and the end of the saccade as completion of 100% of the final duration. Interpolation was done at equally spaced time bins at every 4% from the start to the end of the saccade. This ensured that all saccades of different durations were temporally aligned.

## SUPPLEMENTARY MATERIAL (S2)

### Mean and variance prediction of the dual model

#### Displacement block output

The mean burst generator output from displacement block is obtained by considering the system without any noise signals affecting the system. The expression for the mean signal can then be obtained by taking the expectation of *b*_*nd*_(*t*_*k*_) and can be written as

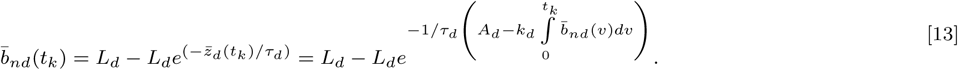

After approximating the integration with the trapezoidal rule, Eq.13 is solved for 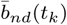. The variance of the displacement block’s output is obtained as

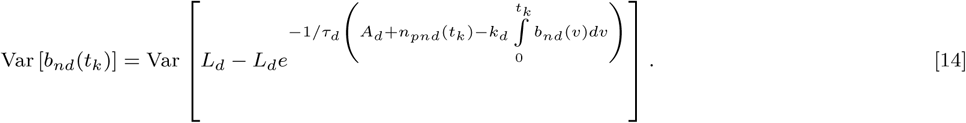

In Eq.14, the terms inside the exponential function are assumed to be independent and hence it can be expanded as

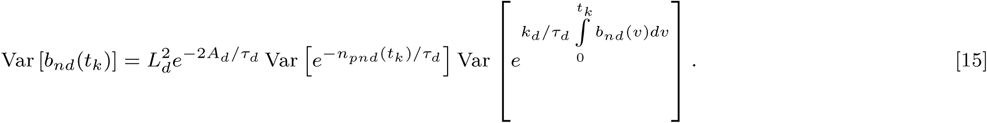

Another assumption is that all random variables under consideration are independent and have a Gaussian distribution. Using the following property of a Gaussian random variable *X* with mean *µ*_*X*_ and variance *σ*_*X*_ ^2^

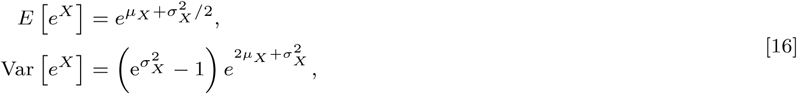

where 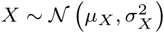. Using this property Eq.15 can be expanded as

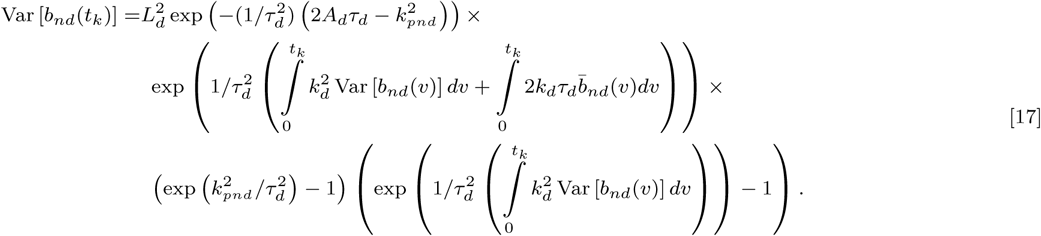

Eq.17 can be solved for Var [*b*_*nd*_(*t*_*k*_)] numerically after approximating the integrals using the trapezoidal rule and using the 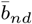 calculated using Eq.13.

#### Velocity block output

Getting the mean and variance of the velocity block output is easier as the burst generator is modeled as a linear function. The mean output is obtained by taking expectation of *b*_*nv*_(*t*_*k*_). It can be written as

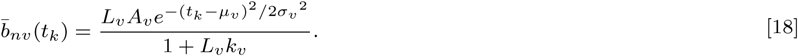

The variance of output of velocity block *b*_*nv*_ (*t*_*k*_) is given by

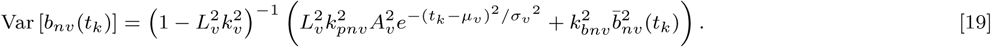

#### Final model output

The required prediction from the model is the mean and variance of angular displacement produced by the oculomotor plant. The mean angular displacement predicted by the model is obtained by using expression of the response of the second order oculomotor plant as

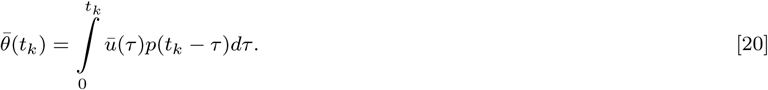

where 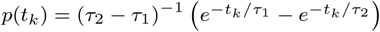. The expression of the mean control signal *ū*(*t*) is the expectation of *u*_*n*_(*t*_*k*_) which is given by

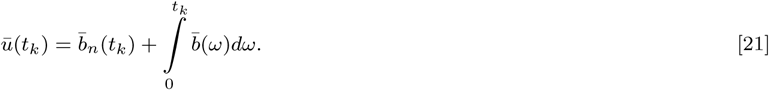

where 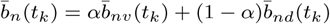. Variance in the angular displacement is given by

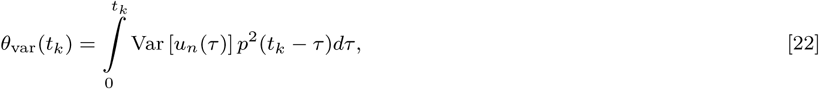

where, the variance in the control signal *u*_*n*_(*t*_*k*_) is obtained by

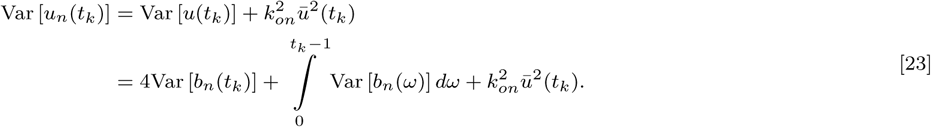

where *b*_*n*_(*t*_*k*_) = *αb*_*nv*_(*t*_*k*_) + (1 − *α*)*b*_*nd*_(*t*_*k*_). The function Var [*b*_*n*_(*t*_*k*_)] can be obtained in terms of Var [*b*_*nd*_(*t*_*k*_)] and Var [*b*_*nv*_(*t*_*k*_)], which are obtained by solving Eq.15 and Eq.19 respectively.

### Reduced stochastic models

Reduced stochastic models are required for obtaining the guess values for parameter estimation in the proposed model as well as for comparison of performance. Hence, reduced stochastic displacement-only and reduced stochastic velocity-only models of saccade generation were created. The reduced stochastic displacement model can be obtained by replacing *b*_*n*_(*t*_*k*_) with *b*_*nd*_(*t*_*k*_) instead of the convex combination. The inter-trial variance and mean profiles of the angular displacement can be derived by following the same procedure as in the dual model. For the reduced stochastic velocity model, *b*_*n*_(*t*_*k*_) needs to be replaced with *b*_*nv*_(*t*_*k*_).

### Model parameter estimation

The model prediction of mean and inter-trial variance profiles of angular displacement is used to form an objective function for the estimation of the free parameters in the model. The model in total has fifteen free parameters. These include nine parameters of the deterministic system which consists of four parameters of the displacement block (*A*_*d*_, *L*_*d*_, *τ*_*d*_, *k*_*d*_) related to the input, burst generation, and internal feedback, five parameters of the velocity block (*A*_*v*_, *µ*_*v*_, *σ*_*v*_, *L*_*v*_, *k*_*v*_) related to input, burst generator, and feedback. Another five parameters related to the noise signal affecting the system includes two noise parameters of the displacement part (*k*_*pnd*_, *k*_*bnd*_), two noise parameters of the velocity part(*k*_*pnv*_, *k*_*bnv*_) and the oculomotor noise parameter (*k*_*on*_). Apart from this, the weightage *α* given to the velocity information is also a free parameter. To find parameters for each subject that fits the observed experimental data of mean and inter-trial variance profile of angular displacement, an objective function in terms of errors is defined as

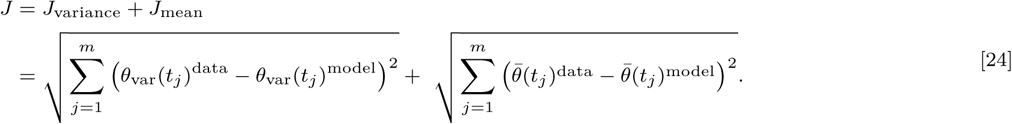

The parameters obtained from the reduced stochastic model fits are used as the guess for the final estimation of all 15 parameters of the stochastic dual model.

## References

1. AA Faisal, LP Selen, DM Wolpert, Noise in the nervous system. Nat. reviews neuroscience 9, 292–303 (2008).

2. CM Harris, DM Wolpert, The main sequence of saccades optimizes speed-accuracy trade-off. Biol. cybernetics 95, 21–29 (2006).

3. RJ Van Beers, Saccadic eye movements minimize the consequences of motor noise. PloS one 3 (2008).

4. CM Harris, On the optimal control of behaviour: a stochastic perspective. J. neuroscience methods 83, 73–88 (1998).

5. H Chen-Harris, WM Joiner, V Ethier, DS Zee, R Shadmehr, Adaptive control of saccades via internal feedback. J. Neurosci. 28, 2804–2813 (2008).

6. T Eggert, FR Robinson, A Straube, Modeling inter-trial variability of saccade trajectories: effects of lesions of the oculomotor part of the fastigial nucleus. PLoS computational biology 12 (2016).

7. D Robinson, The use of control systems analysis in the neurophysiology of eye movements. Annu. review neuroscience 4, 463–503 (1981).

8. CA Scudder, A new local feedback model of the saccadic burst generator. J. neurophysiology 59, 1455–1475 (1988).

9. H Goossens, AJ van Opstal, Optimal control of saccades by spatial-temporal activity patterns in the monkey superior colliculus. PLoS Comput. Biol. 8 (2012).

10. R Jürgens, W Becker, H Kornhuber, Natural and drug-induced variations of velocity and duration of human saccadic eye movements: evidence for a control of the neural pulse generator by local feedback. Biol. cybernetics 39, 87–96 (1981).

11. DA Robinson, Models of the saccadic eye movement control system. Kybernetik 14, 71–83 (1973).

12. I Smalianchuk, UK Jagadisan, NJ Gandhi, Instantaneous midbrain control of saccade velocity. J. Neurosci. 38, 10156–10167 (2018).

13. A Van Opstal, J Van Gisbergen, J Eggermont, Reconstruction of neural control signals for saccades based on an inverse method. Vis. research 25, 789–801 (1985).

14. CM Harris, DM Wolpert, Signal-dependent noise determines motor planning. Nature 394, 780–784 (1998).

15. RJ van Beers, The sources of variability in saccadic eye movements. J. Neurosci. 27, 8757–8770 (2007).

16. JB Smeets, IT Hooge, Nature of variability in saccades. J. neurophysiology 90, 12–20 (2003).

17. E Bollen, et al., Variability of the main sequence. Investig. ophthalmology & visual science 34, 3700–3704 (1993).

18. D Schmidt, L Abel, L DellOsso, R Daroff, Saccadic velocity characteristics-intrinsic variability and fatigue. Aviat. space, environmental medicine 50, 393–395 (1979).

19. KE Jones, AFdC Hamilton, DM Wolpert, Sources of signal-dependent noise during isometric force production. J. neurophysiology 88, 1533–1544 (2002).

20. DM Waitzman, TP Ma, LM Optican, RH Wurtz, Superior colliculus neurons mediate the dynamic characteristics of saccades. J. Neurophysiol. 66, 1716–1737 (1991).

21. TR Reppert, M Servant, RP Heitz, JD Schall, Neural mechanisms of speed-accuracy tradeoff of visual search: saccade vigor, the origin of targeting errors, and comparison of the superior colliculus and frontal eye field. J. neurophysiology 120, 372–384 (2018).

22. DL Sparks, L. Mays, Signal transformations required for the generation of saccadic eye movements. Annu. review neuroscience 13, 309–336 (1990).

23. A Berthoz, A Grantyn, J Droulez, Some collicular efferent neurons code saccadic eye velocity. Neurosci. letters 72, 289–294 (1986).

24. K Arai, S Das, EL Keller, E Aiyoshi, A distributed model of the saccade system: simulations of temporally perturbed saccades using position and velocity feedback. Neural Networks 12, 1359–1375 (1999).

25. A Van Opstal, J Van Gisbergen, Scatter in the metrics of saccades and properties of the collicular motor map. Vis. research 29, 1183–1196 (1989).

